# On Global Brain Reconfiguration after Local Manipulations

**DOI:** 10.1101/2023.09.08.556815

**Authors:** Giovanni Rabuffo, Houefa-Armelle Lokossou, Zengmin Li, Abolfazl Ziaee-Mehr, Meysam Hashemi, Pascale P Quilichini, Antoine Ghestem, Ouafae Arab, Monique Esclapez, Parul Verma, Ashish Raj, Alessandro Gozzi, Pierpaolo Sorrentino, Kai-Hsiang Chuang, Teodora-Adriana Perles-Barbacaru, Angèele Viola, Viktor K. Jirsa, Christophe Bernard

**Affiliations:** Institut de Neurosciences des Systèms, Aix Marseille University, UMR 1106 INSERM, Marseille, France; Center for Magnetic Resonance in Biology and Medicine, Aix-Marseille University, UMR 7339 CNRS, Marseille, France; Queensland Brain Institute, The University of Queensland, Brisbane, Australia; University of California, UCSF, San Francisco, USA; Functional Neuroimaging Laboratory, Istituto Italiano di Tecnologia, Rovereto, Italy

**Keywords:** Mouse fMRI, Resting-state, Thalamic lesion, Chemogenetic silencing, Brain models, Inference, Multiscale mechanisms

## Abstract

Understanding how localized brain interventions translate into whole-brain dynamics is crucial for deciphering neural function and tailoring therapeutic strategies. Combining mouse experimental datasets of focal interventions (thalamic lesion and chemogenetic silencing of cortical hubs), we demonstrate both local and global effects. Using whole-brain simulations of experimental data, we not only confirm the distributed nature of local manipulations but also offer mechanistic insights into these processes. Our simulations predict specific alterations in firing rates and spectral characteristics across specific brain networks, leading to structured changes in functional connectivity patterns. Some of these predictions have been empirically validated. Notably, the affected brain subnetworks—and their resultant ‘signatures’ of change—are contingent on the original intervention site, suggesting a method to accurately localize the source of alteration. Our results provide a general framework for interpreting localized intervention effects, offering insights that could refine clinical interventions for focal brain disorders by enabling targeted circuit-level neuromodulation strategies.

## 1 Introduction

A choice method to obtain insights into brain function and dysfunction is up- or down-regulating neuronal activity and studying the brain’s response to such perturbations. This allows for establishing cause-effect relationships and proposing mechanisms. For example, optogenetics and chemogenetics have become essential for delineating the roles of specific neurons within distinct brain regions, thus allowing researchers to map functions and behaviors to cellular activities [1, 2]. However, the inference from such mappings to assert that certain neurons govern specific behaviors is not straightforward. The brain, as a complex system, underscored by emergent phenomena that manifest at a systemic level beyond the sum of its parts, questions such straightforward attribution [3]. Neurons firing in correlation with a given information processing/action tend to be widely distributed throughout the brain [4]. Thus, controlling neuronal activity in one region may not be sufficient to affect information processing/action because of the distributed nature of brain processing.

Another argument supporting the idea that inference from such mappings is not straightforward stems from the fact that affecting a few components in a complex system may drastically change its behavior. The activity of a few neurons has the potential to reshape the system-level organization [5]. This concept is exemplified by the phenomenon of focal diaschisis, where localized stimulation or lesions induce both proximal and distal effects, altering activity in regions far removed from the intervention site [6, 7]. Studies in humans and animal models have documented such distant responses to focal lesions, profoundly affecting functional connectivity and organization [6, 8–12]. Moreover, targeted manipulations of neuronal activity have been shown to modulate functional connectivity, with both enhancements and reductions observed depending on whether the neurons were stimulated or inhibited [13–18]. Nevertheless, the mechanisms by which local interventions result in global brain activity changes remain elusive.

The implications of these findings could be far-reaching, particularly for the development of targeted therapies such as deep brain stimulation and transcranial magnetic stimulation. The current empirical basis for these interventions stands to be significantly advanced by a deeper understanding of how localized neuronal modulation translates to whole-brain dynamics. Bridging this knowledge gap could transform therapeutic strategies, fostering more refined and theoretically informed approaches.

Here, we study the global repercussions of localized neuronal interventions by examining two independent functional magnetic resonance imaging (fMRI) datasets from mice. These datasets provide a before-and-after snapshot of whole brain activity following specific site silencing — one targeting thalamic regions and the other cortical hubs. We leverage a whole mouse brain model [19, 20] and simulation-based inference (SBI) [21–23] to virtualize the mouse brain and study the effects of focal silencing at the physiological and functional levels in silico.

## 2 Results

We analyzed two independent datasets (Fig.1.A, Suppl. Fig.1) to investigate the effects of inactivating a focal area of the brain on whole-brain dynamics. In a ‘Lesion’ dataset, we used an irreversible lesion of thalamic nuclei by excitotoxic neurodegeneration produced by local administration of N-methyl-D-aspartate (NMDA) injection (Th; 6 C57BL6 mice). In this group, Blood-oxygen-level-dependent (BOLD) acquisitions were performed in ‘resting-state’ fMRI (rs-fMRI) under light anesthesia before and 6 weeks after the lesion (Suppl. Fig.1, histology). In a ‘DREADD’ dataset, C57Bl/6J mice were recorded under light anesthesia, before and 30 minutes after clozapineNoxide (CNO) injection causing DREADD-mediated pan-neuronal inhibition of either the anterior cingulate area (ACA; 7 mice) or the retrosplenial cortex (RSC; 14 mice). A control group (CTRL; 8 mice) underwent the same experimental pipeline as the RSC group, but the mice were not expressing DREADD (Suppl. Fig.1, experimental figures).

**Fig. 1.**
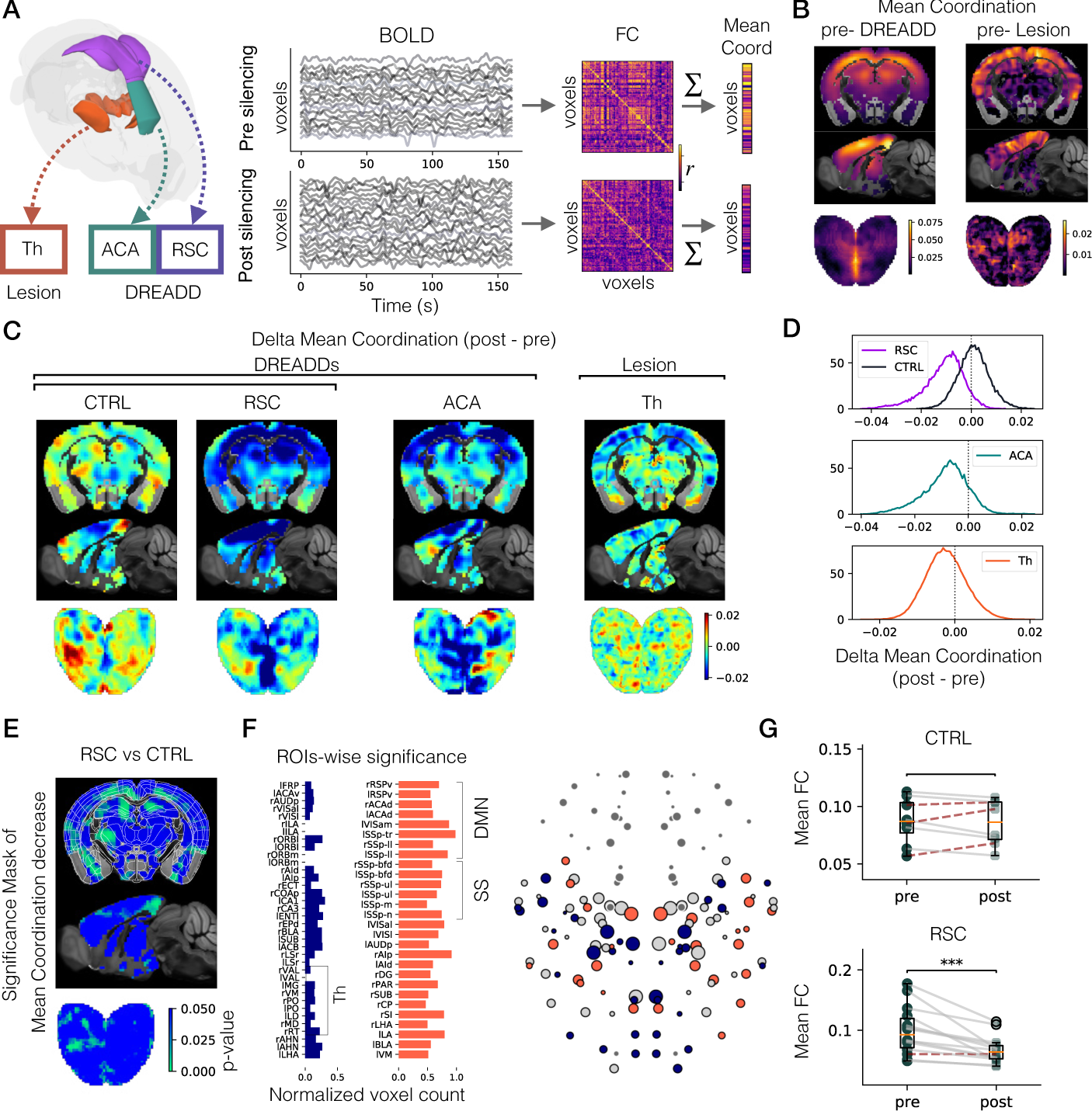
Focal silencing induces functional connectivity reconfiguration: A) We analyzed two independent datasets where mice were recorded before and after silencing a brain region. Silencing was obtained by either lesion of a few Thalamic (Th) nuclei, or by chemogenetic (DREADD) inhibition of the anterior cingulate area (ACA) or the retrosplenial (RSC; dorsal and ventral) cortices. For pre- and post-silencing data we extracted the voxel-level FC and the mean Coordination (i.e., the FC degree), defined as the average correlation of the voxel with the rest of the voxels. B) Mean Coordination obtained by averaging across prelesion data and all pre-DREADD data. Note that the mean Coordination decreases along a dorsoventral axis and is higher in cortical regions. C) The Delta (post-pre) Mean Coordination (DMC) plotted for all the groups reveals that, after the focal silencing, a distributed decrease of mean Coordination is observed (dark blue) in all groups. The control group remains globally similar between post and pre (CTRL; sham of the RSC). D) The distribution of the DMC values in panel (C) is shifted to the left and skewed in all groups but not in the CTRL, which indicates an overall decrease in mean Coordination after focal silencing. (E) For the statistical significance of the mean Coordination decrease, we performed the Mann-Whitney U test in each voxel, comparing mice in the RSC group to mice in the CTRL group (green values indicate significance *p* < 0.05). F) Regions of interest (ROIs) with an unexpected percentage of decreased and increased voxels compared to null models. Regions showing an unexpected decrease (in blue) include most Thalamic nuclei, while those showing an unexpected increase (in red) include regions within the Default Mode Network (DMN) and the Somatosensory (SS) network, among others. G) RSC silencing resulted in decreased mean FC in 13 mice (continuous grey lines) and an increase in 1 mouse (dashed red lines). In the CTRL group, 5 mice decreased and 3 increased post-CNO injection. The mean FC was significantly decreased in the RSC group only (Wilcoxon signed-rank test *p* < 0.001).

We first compared voxel-level Functional Connectivity (FC), defined as Pearson’s correlation over all voxel pairs, before and after focal silencing (Fig.1.A). Figure 1.B shows, for each voxel the mean of the FC matrix across one of the two axes, averaged across all pre-lesion (left) and pre-DREADD (right) mice. This measure is sometimes referred to as ’weighted degree centrality’ [24] or ’global connectivity’ [25] since the FC degree corresponds to the average correlation between a voxel and the rest of the network. For the sake of consistency with the forthcoming analyses, we will use the term ‘mean Coordination’ to refer to this measure (see also Fig.2.A and subsection 2.2). Upon visual inspection, these brain plots reveal that cortical voxels are well coordinated with the whole-system activity, while the mean Coordination gradually decreases in a gradient towards subcortical structures. These results were consistent across datasets, despite the different spatial and temporal resolutions (see Methods). In the DREADD groups, for which we have more mice with higher temporal resolution (2000 time points), the average pre-DREADD mouse data revealed a strong coordinated core (in yellow) along the cortical midline structures, with a peak in the deep cortical layers and the dentate gyrus. These regions roughly correspond to what was previously identified as the mouse default mode network (DMN; [26]). These results show that, as previously reported[25], mouse rs-fMRI is organized in functional networks, with specific subsets of voxels that are highly coordinated to whole-brain network fluctuations

**Fig. 2.**
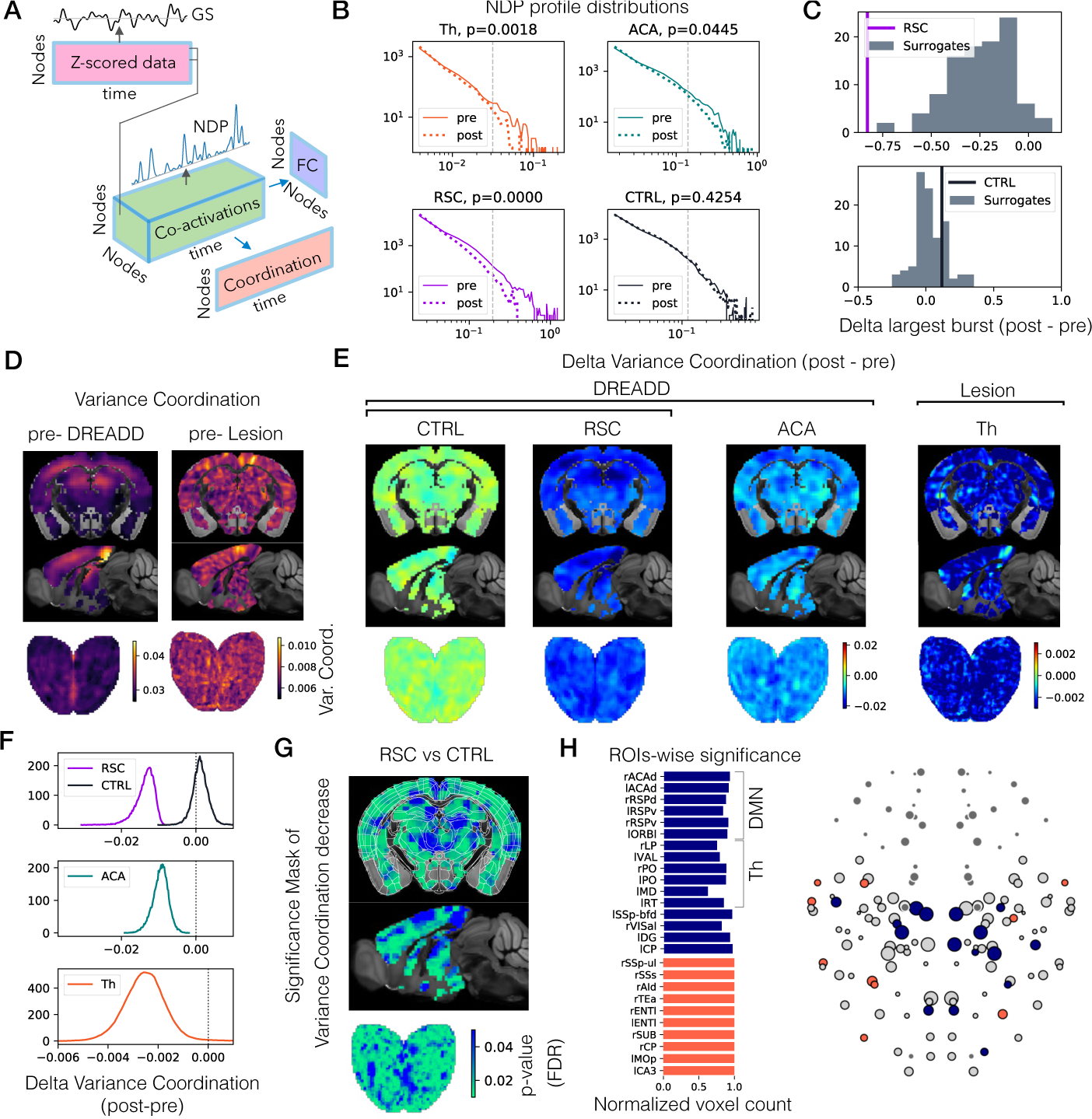
Dynamic origin of FC decrease after focal lesion: A) Illustration of the analytic link between z-scored brain signals, edge co-activations, dynamic network profile (DNP), FC, and the Coordination measure. Each arrow represents the correlation along the parallel axis. Here the nodes correspond to voxels or ROIs. B) The pre-silencing NDP distribution (continuous lines) is fat-tailed, indicating the presence of large-scale bursts of co-activations in the data. After focal silencing (dashed lines) the NDP decreases, showing an impaired capability of bursting. This effect was significant in RSC, ACA, and Th groups, but not in the CTRL group (Kolmogorov–Smirnov test performed for values above the dashed vertical lines; see p-values in the figure). C) The difference in the size of the largest burst between post and pre-silencing i.e., the largest value of the NDP is shown in purple and black for the RSC and CTRL groups, respectively. Surrogates (grey distributions) cannot explain the largest NDP bursts in the RSC. D-H) As panels B-F in Figure 1, for the Coordination variance instead of the mean Coordination. These results reveal even more widespread effects of focal silencing than those observed in Figure 1, which were based on static FC

### 2.1 Widespread decrease of functional connectivity after focal region silencing

To probe the impact of focal region silencing on brain-wide measure of FC, for each mouse, we evaluated the difference between the mean Coordination post- and pre-silencing (Delta Mean Coordination; DMC). Analyses of Th-, ACA-, and RSC-manipulated mice, revealed that after lesion/silencing, the mean Coordination patterns were decreased beyond the local neighborhood of the target regions (Fig.1.C). FC measurements vary in test-retest conditions in the same mouse [27], which explains positive and negative values in the CTRL group (Fig.1.C). However, the DMC observed in the CTRL group was much smaller than in the RSC group (Fig.1.D, Kolmogorov-Smirnov statistics *KS* = 0.62, *p* < 0.001).

The DMC distribution showed a leftward shift for Th, ACA, and RSC groups but not for the CTRL, showing that a general decrease of whole-brain FC occurs after silencing (Fig.1.D; mean values: *µ_th_* = −0.0026*, µ_ACA_* = −0.0087*, µ_RSC_* = −0.0113*, µ_CTRL_* = 0.0007). Despite this general decrease, the mean Coordination increased in a subset of voxels (positive DMC values) in all groups, indicating that FC also increased along some pathways after focal silencing. We also noted that the DMC distribution of the RSC group was skewed to the left, unlike the CTRL group, which was more symmetric (Fisher-Pearson skewness coefficient *g_RSC_* = −0.72*, g_CTRL_* = 0.10). The same trend was evident for ACA (*g_ACA_* = −0.38). This indicates that a subset of voxels is more prominently affected by chemogenetic silencing than others. In contrast to RSC and ACA, the Th group showed, to a lesser extent, some skewness to the left (*g_Th_* = 0.14).

We then identified the voxels most prominently affected by the chemogenetic silencing of RSC, for which we have the control group (CTRL). For each voxel, we compared with a Mann-Whitney U test the difference in mean Coordination values measured before and after chemogenetic silencing. In Figure 1.E, we plot the significance values for each voxel. Importantly, the significance mask exhibited high spatial structure, demonstrating a topography of the voxels that significantly decreased after the RSC silencing. To interpret this result, we grouped the voxels according to the Allen Atlas mouse brain parcellation [28] with *N* = 148 regions of interest (ROIs; see Table 1) and computed the percentage of voxels significantly affected by the silencing within each brain region. This percentage was compared against a benchmark by randomly shuffling the significance mask. In Figure 1.F, we show in blue the ROIs with an unexpectedly low number of significantly decreased voxels (i.e., ROIs that were expected to have more significant voxels according to the randomized masks), including a distributed network of regions and most thalamic nuclei. The ROIs with a higher-than-expected percentage of significantly decreased voxels (in red), include DMN and somatosensory (SS) ROIs, among other regions.

Finally, we show that a chemogenetic-induced decrease in correlation is also observable at the level of ROIs-reconstructed signals (see Appendix A). To this end, we first averaged the activities of all the voxels belonging to each region, which defined region-specific (order of ∼ 1−10*mm*^3^) BOLD time series. Adjacency (FC) matrices were computed for each mouse before and after region silencing. Figure 1.G shows the average FC for each mouse of the RSC and CTRL groups, which significantly decreased only in the RSC group. Together, these results show that silencing specific brain regions results in widespread but structured reorganization of large-scale FC at rest. Because FC provides a static vision by averaging activity over the scanning time, we next performed a more detailed analysis focusing on brain dynamics at the timescale of the BOLD signal.

### 2.2 Changes in whole brain dynamics after focal region silencing

At rest, brain dynamics is characterized by the occurrence of intermittent bursts of activity [29–31]. During such bursts, many (sometimes very distant) brain regions display sudden co-fluctuations, giving rise to system-level correlations [32]. We thus hypothesized that the observed disruption of system-level correlations following focal region silencing (Fig.1) would be associated with a reduced capability of the network to generate widespread coordinated bursting. To test this hypothesis, we use an edge-wise analysis [29], whereby regions (or voxels) are taken as network nodes. The element-wise product of any two z-scored regional signals *Z_i_*(*t*) and *Z_j_*(*t*) define an edge co-fluctuation time series *E_ij_*(*t*) = *Z_i_*(*t*) · *Z_j_*(*t*) for regions *i* and *j* (Fig.2.A)

The edge co-activation matrix can be considered as a time-resolved FC at the timescale of the BOLD signal since the Pearson’s correlation coefficient between node *i* and *j* is defined as the time average of their co-fluctuations i.e., *r* = ⟨*E_ij_*(*t*)⟩*_t_*. In other words–as sketched in Figure 2.A–the FC matrix (in purple) corresponds to the time average of the co-activation matrices (in green). Averaging the co-activation signals across all network edges defines the network dynamic profile *NDP* (*t*) = ⟨*E_ij_*(*t*)⟩*_ij_*. Such profile (corresponding to the square of the global signal *GS*(*t*) = ⟨*Z_i_*(*t*)⟩*_i_*; see Methods) follows a fat-tailed distribution (Fig.2.B). Rare high-amplitude events in the NDP correspond to strong simultaneous co-fluctuations of a large set of edges.

We compared the NDP profiles (pooled together for all mice within each group) before and after the regional silencing (Fig.2.B). The NDP distribution of the Th, ACA, and RSC groups exhibited a faster decrease in post-silencing recordings than in pre-silencing ones (the p-values of the Kolmogorov-Smirnov statistic in Fig.2 are evaluated considering bins above the dashed gray line). This result aligns with our hypothesis that the observed FC decrease caused by focal silencing is associated with a decrease in spontaneous network bursting. Importantly, this decrease does not occur in the CTRL group (Fig.2.B).

The decrease in large network bursts observed in the RSC may be a direct consequence of the decrease in FC identified above (Fig.1). Alternatively, it may reflect a genuine dynamical impairment. To disambiguate this dichotomy, we compared the NDP distribution to 100 surrogate datasets obtained from phase-randomization (cross-spectrum preserved) of the voxel signals (see Methods). This kind of surrogate data maintains the static correlation between voxel pairs (i.e., the static FC) so that each surrogate shows the same decrease of mean Coordination displayed in the panels in Figure 1. In Figure 2.C, we show that the maximum burst size observed in the RSC data (purple vertical line) is significantly lower than the maximum value expected from the surrogate NDPs (gray distribution). The same result does not hold for the CTRL group. We conclude that the decrease in static FC observed in our data does not fully explain the absence of large-scale bursts after the silencing of RSC. Thus, focal silencing impairs spontaneous network bursting.

### 2.3 Altered local to global coordination dynamics

The change in spontaneous network bursting may be due to a disruption in the way some local nodes are coordinated with the rest of the brain. To test this hypothesis, we defined a metric of local-to-global Coordination, which characterizes the degree of participation of voxel *i* (or network node *i*) to the global signal fluctuations in a time-resolved way. Coordination is defined as the element-wise product *C_i_*(*t*) = *Z_i_*(*t*) · *GS*(*t*) of the voxel (standardized) activity *Z_i_*(*t*) and the global signal *GS*(*t*). The Coordination corresponds to the average co-fluctuations of all the edges *ij* hinging on voxel *i* i.e., *C_i_*(*t*) = ⟨*E_ij_*(*t*)⟩*_j_* (as illustrated in Fig.2.A).

For a given voxel, strong Coordination corresponds to moments when the voxel is co-fluctuating with the rest of the network, whereas weak Coordination corresponds to moments when the voxel is disjoint from the rest of the brain (or when the global signal is close to zero). In Figure 2.D-H, we show that the variance of the Coordination displays a precise, non-random topography and its alteration due to focal lesion/silencing is reminiscent of the voxel-level differences described in Figure 1.B-E. The variance of the Coordination represents the ability of a region to dynamically fluctuate in and out of coordination with the rest of the network. Fluctuations are considerably reduced after the chemogenetic silencing of RSC, while they are not affected in the CTRL group (Fig.2.G-H). The analysis of the Coordination variance revealed stronger and even more widespread effects of focal silencing, as compared to the mean Coordination for all RSC, ACA, and Th groups. For example, comparing Figure 1.D to Figure 2.F, one can observe that the distributions in the latter are further left to zero, indicating a net decrease in voxel dynamics. The voxelwise significance test in Figure 2.G highlights widespread significant changes in Coordination variance that, differently from Figure 1.D, survive the FDR test. ROIs-level analysis revealed that fewer voxels than expected decreased in Coordination in several DMN and thalamic regions (Fig.2.H).

Collectively, these results demonstrate a clear reduction in static FC and dynamic network properties following either the chemogenetic inhibition of cortical hubs (RSC and ACA) [33] or the lesion of thalamic (Th) nuclei. To identify underlying mechanisms, in particular at the electrophysiological level, we must turn to computational approaches, specifically whole-brain modeling. Such models offer a unique and powerful framework to simulate and analyze brain-wide interactions to understand emergent behaviors and make testable non-trivial predictions. In addition, the computational approach may help us understand the origin of the discrepancy between our work and previous ones, which suggests fMRI over-connectivity after focal chemogenetic silencing in the mouse [16] and non-human primates [17].

### 2.4 Virtual Mouse Brain simulation

We simulated whole-brain activity using The Virtual Mouse Brain (TVMB) pipeline [19, 20] (see Methods). We focused on the RSC group and we built an in silico model of the pre-DREADD fMRI activity, that reproduced the average data features observed in the RSC mice before silencing. Simulated data (Fig.3.A) includes fast neuroelectric activity (e.g., average firing rate dynamics; 1000Hz sampling rate) as well as slow BOLD activity (0.5Hz sampling rate), which can be directly compared to empirical observations. The simulated activities depend on the choice of local and global model parameters. The main parameters for our simulations are the local node excitability *η* and the global coupling *G*. The latter measures the impact of structural connectivity on local dynamics: for *G* = 0 the regions are effectively decoupled while increasing *G* the regions are more driven by their mutual coupling. Brain heterogeneity was modeled by associating different excitability values to nodes belonging to different mouse resting-state networks identified according to a common classification (as in Suppl. Fig.2): Default Mode Network (DMN), Lateral Cortical Network (LCN), Visual Network (Vis), Basal-Frontal Network (BF), Hippocampal Formation (HPF), and Thalamus (TH). In total, our model has seven free parameters (*G, η_DMN_, ηV_is_, η_LCN_, η_BF_, η_HPF_, η_Th_*).

**Fig. 3.**
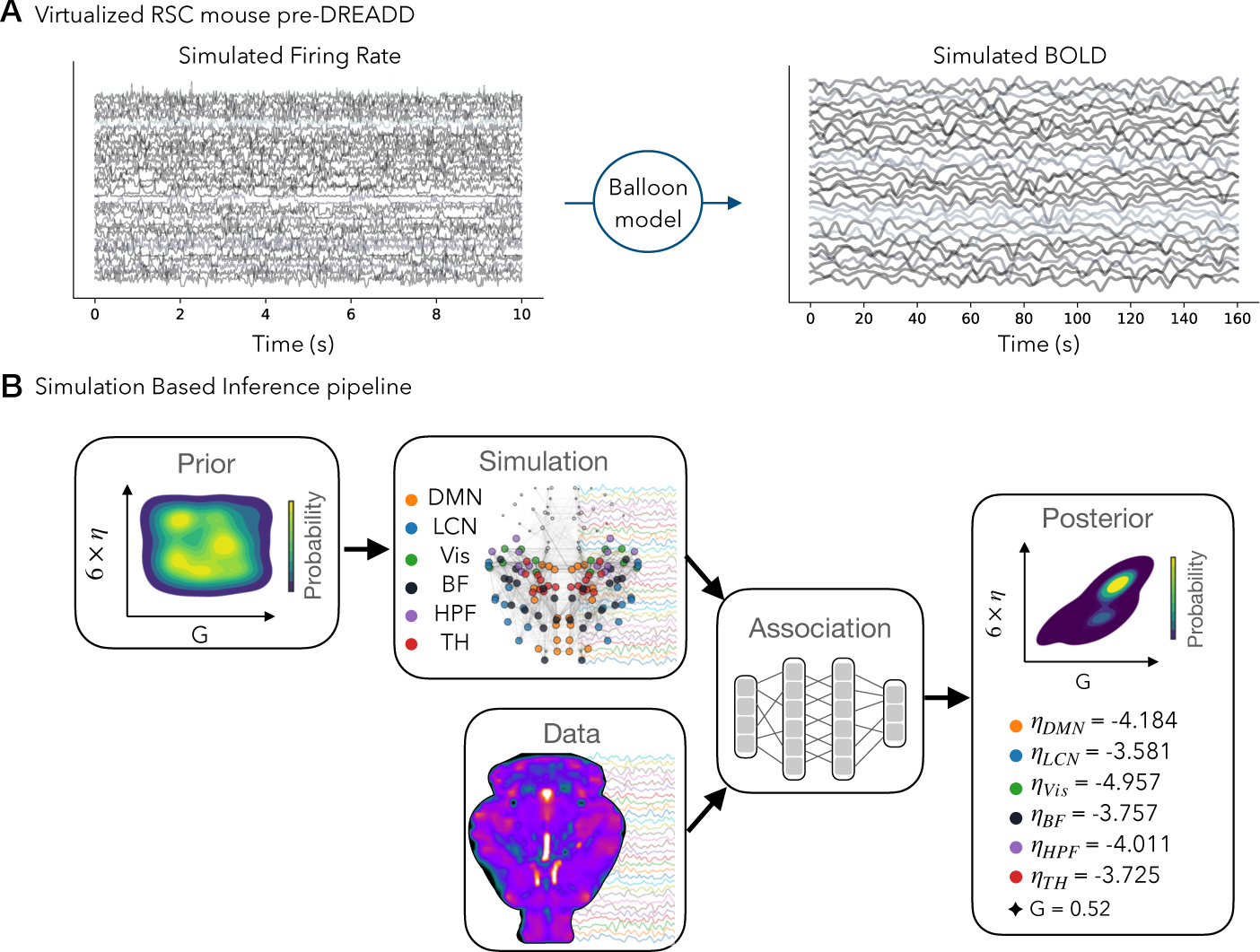
Virtual Mouse Brain and Simulation-Based Inference: A) The Virtual Mouse Brain pipeline allows the generation of fast neuroelectric (firing rate, left) and slow hemodynamic (BOLD, right) activities in silico. Here we simulated realistic brain activity for the average RSC mouse pre-DREADD. To closely match the empirical data features, we need to infer the local and global model parameters. B) The best configuration of model parameters was found by a Simulation-Based Inference (SBI) approach. In short, uniform prior probability distributions of global coupling (G) and regional excitability (6 *η*values) were given as the pipeline input. Then, a large number of simulations were generated using random parameters taken from the prior distribution. In each simulation, data features were extracted for probabilistic inference against empirical data. Deep neural density estimators were trained to learn the association between simulations and data. Finally, the SBI outputs a posterior probability distribution quantifying the best working point where simulations match observations, with the associated uncertainty in the estimation.

To simulate realistic empirical data, it is required the identification of the sets of free parameters that most closely reproduce empirical pre-DREADD data in silico. To this aim, we used a simulation-based inference (SBI) work-flow, which is well suited for exploring such a large parameter space (Fig.3.B). SBI takes as the input a prior probability distribution of the parameters’ values, compares simulated and observed data features across several parameter values, and uses a probabilistic deep learning model to output a posterior distribution peaked at a configuration of parameters where the simulations best match the empirical data [21] (see Methods). Importantly, SBI allows for inference on multimodal distributions and parameter degeneracy, as compared to classical variational inference used in Dynamic Causal Modeling [34, 35].

### 2.5 Connectional diaschisis in silico: BOLD analysis

Having performed parameter estimation to reproduce pre-DREADD activity in the RSC group, we reasoned that if our simulations were to bear any physiological relevance, the focal silencing in silico should produce local and large-scale effects without further parameter manipulation. We first used the working point parameters listed in Fig.3.B, running *N* = 264 baseline (Base) simulations by changing the initial conditions. We then performed two types of manipulations, manually increasing or decreasing by ∼ 10% the excitability of the (left and right, ventral and dorsal) RSC without changing any other parameter value. We compared the three arrays of simulations whose settings only differed in the excitability value of the retrosplenial cortex: *η_RSC_* = *η_DMN_* = −4.184 in the simulated pre-silencing (Base), higher excitability *η_RSC_* = −3.8 in the simulated focal excitation (Exc), and lower excitability *η_RSC_* = −4.6 in the simulated focal inhibition (Inh). Despite the local nature of the manipulations, the average BOLD-FC significantly changes at the whole brain level (Fig.4.A, left). If RSC excitability is decreased, FC increases (Fig.4.A, left), in line with experimental results reporting overconnectivity post-DREADD inhibition [16]. Conversely, FC decreases if RSC excitability increases (Fig.4.A, left). In agreement with the results shown in Figure 2.A-B, a decrease in FC was associated with a decreased NDP, and conversely, an increase in FC was associated with increased bursting i.e., larger events in the NDP distribution (Fig.4.A, right). Correlation between Functional Connectivity and Structural Connectivity (SC) has been identified as a relevant marker of brain function, whereby FC-SC correlations deviate from healthy values in several pathologies [36, 37]. Since we use the Allen connectome to build TVMB, we can calculate FC-SC correlations after modifying neuronal activity. We found that increased (decreased) FC was associated with a greater (lower) correlation between FC and SC (Fig.4.B). Finally, we compared the simulated data with the empirical BOLD data reconstructed at the ROI level. The delta FCs–defined as the difference between the FC in the case of focal excitation or inhibition, minus the baseline FC*_base_* (Fig. 4.C, top)–show that modulating retrosplenial excitability results in a strong effect over DMN and Visual networks and in their connections to the Th network. These results confirm that focal brain manipulations result in large-scale effects: focal diaschisis.

**Fig. 4.**
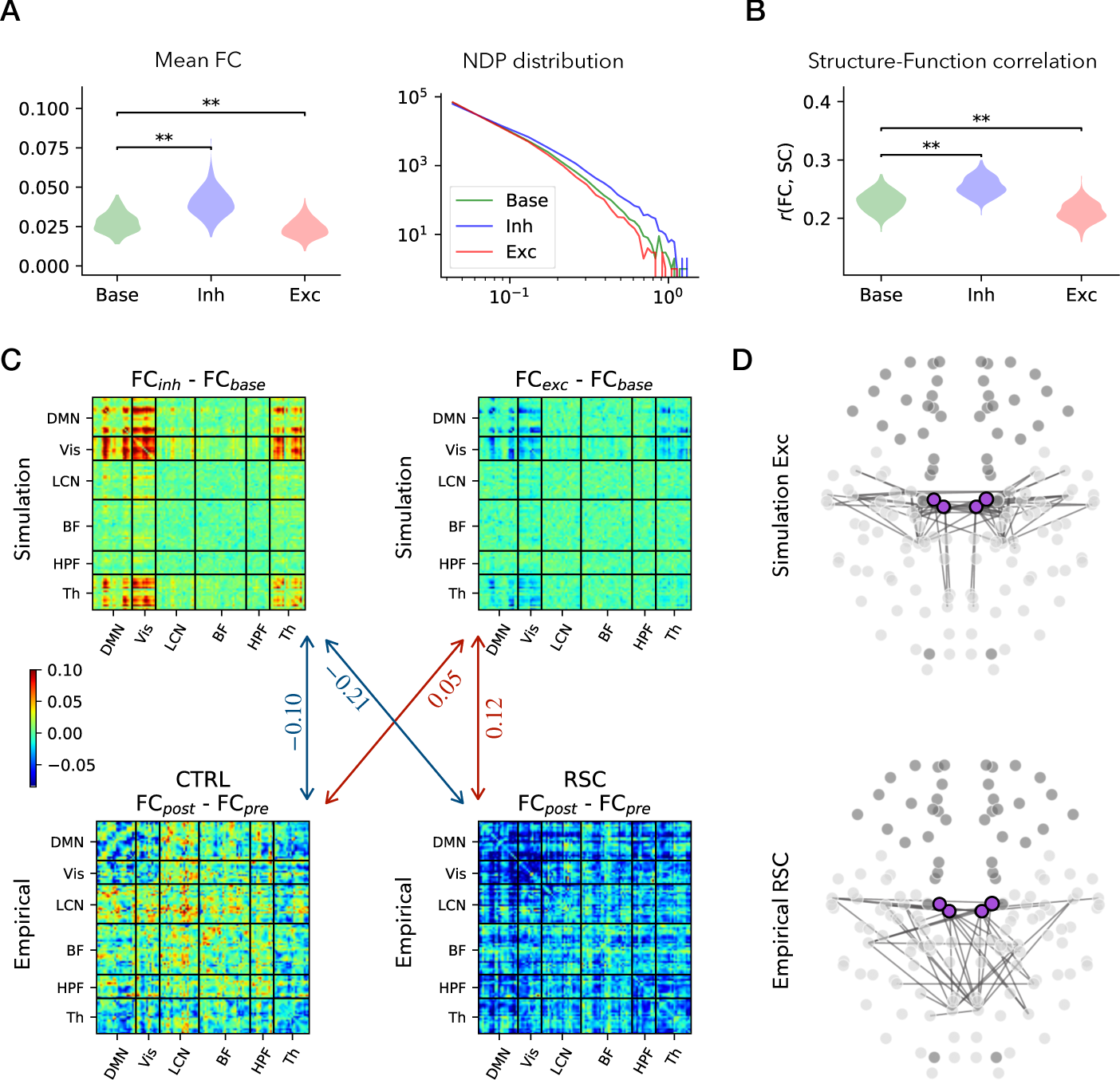
Connectivity changes after focal excitability modulation in silico: A) The average FC increased after focal inhibition, while it decreased after focal excitation (Wilcoxon test: *p* < 0.001 (*), *p* < 0.0001 (**)). The NDP distribution changed accordingly, with less large bursts associated to decreased FC, and vice versa. B) The Pearson correlation between FC and the Allen SC (used as the structural connectome for the simulations) increased after focal inhibition, while it decreased after focal excitation. C) The simulated FC difference (Exc-Base, or Inh-Base; top row) and the observed FC differences (post-pre; bottom row) for the CTRL and RSC groups. The numbers indicate the Spearman’s *r*s between two matrices linked by the associated arrow (*p* < 0.001). D) The edges that decreased the most in terms of correlation after simulated RSC excitation (top), and the edges with the largest correlation decrease in the observed RSC group (bottom; decrease larger than any edge in CTRL). Purple nodes mark the left/right, ventral/dorsal RSC.

With minimal parameter tuning—restricted solely to those necessary for replicating baseline neural activity—our computational approach produced non-trivial results. It exemplifies an emergent property that is only discernible within the context of the whole brain dynamics. When the activity of a single brain region is altered, the repercussions resonate throughout the brain. However, simulations show that increasing excitability results in decreased FC, whilst inhibiting the RSC decreased FC experimentally (Fig. 1.C). We thus predicted that DREADD inhibition and lesion could result in an unexpected increase of excitability in the targeted ROIs. The general decrease in delta FC observed for RSC DREADD inhibition (Fig.4.C, bottom right) significantly correlates with the decrease observed for simulated focal excitation (Sperman’s *r_s_* = 0.12*, p* < 0.001). The delta FC of the CTRL group shows a weaker correlation (*r_s_* = 0.05*, p* < 0.001). Both delta FC of RSC and CTRL groups anticorrelate with the simulated delta FC in the case of focal inhibition (*r_s_* = −0.21*, p* < 0.001, and *r_s_* = −0.1*, p* < 0.001, respectively).

Despite these correlation values being moderate, they are highly significant, and correlations in the RSC group are always higher than in the CTRL group. Furthermore, the inhibited and excited values of the *η_RSC_* parameter were not fine-tuned to maximize the correlation values, which was beyond the illustrative scope of this analysis. These results support the theoretical prediction that in our experiments focal silencing results in an overall increase of excitability in the target region. For a visual comparison of the altered topography after simulated focal excitation, we plot the 1.5 percentile of the strongest decaying long-range correlations (Fig.4.D, top), which can be compared to the strongest decaying edges observed in the RSC (Fig.4.D, bottom).

These simulations demonstrate connectional diaschisis in silico, allowing us to interpret experimental data, hypothesize the effects of experimental manipulations over cell excitability, and predict the effects of focal excitation and inhibition. After estimating the parameters that enabled to reproduce baseline conditions, we did not perform any further parameter adjustments to fit theoretical and experimental data. Remarkably, our whole-brain simulations produced large-scale effects like those found experimentally and generated predictions that we validated experimentally.

### 2.6 Focal diaschisis in silico: Local Field Potential analysis

Brain simulations also allowed us to investigate the neuroelectric activity underlying the BOLD dynamics in each brain region, an analysis that cannot be performed experimentally. Figure 5.A shows the effects of the modulation (inhibition or excitation) of RSC excitability in the left dorsal RSC (dRSC). Power spectral analysis revealed that inhibition of RSC results in increased slow frequencies and decreased fast frequencies (Fig.5.B), a prediction of the model already validated experimentally [16]. The opposite result is predicted to occur (decreased slow frequencies and increased fast frequencies) when the excitability of the RSC is increased (Fig.5.B).

**Fig. 5.**
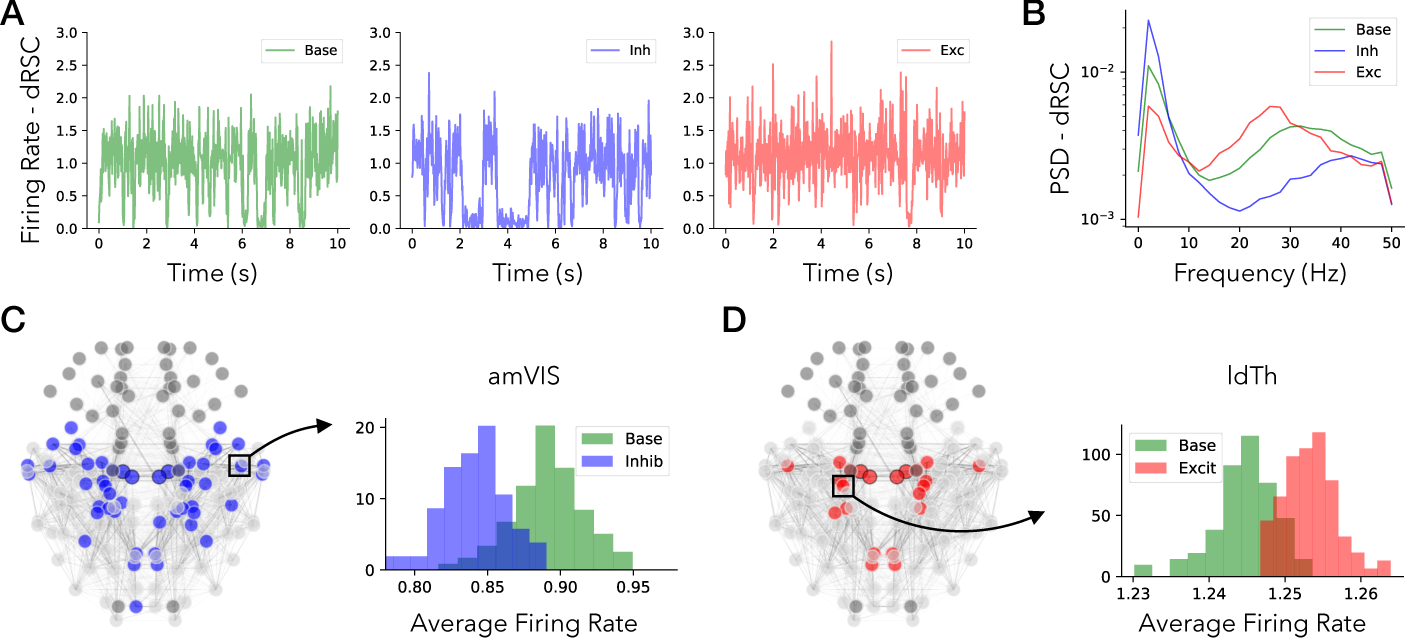
Diaschisis in silico: A) Firing rate activity for the dorsal RSC (dRSC) in Base, Inh, and Exc simulations. B) The PSD for dRSC shows opposite changes after focal inhibition or excitation. C) RSC inhibition leads to a decreased mean firing rate in a distributed network of nodes, such as the anteromedial visual area (amVIS). D) RSC excitation leads to an increased mean firing rate across a distributed subset of ROIs, such as the laterodorsal nucleus of the Thalamus (ldTh).

Studying the simulated fast firing rate dynamics, we further show that the activity also changes in brain regions away from the silenced region, further demonstrating diaschisis (Fig.5.C-D). In particular, the inhibition of RSC resulted in a significant firing rate decrease in a subnetwork of ROIs (blue nodes in Figure 5.C). An example of firing rate distributions before and after RSC inhibition is shown for the left anteromedial visual cortex (amVIS; Fig.5.C, right). Conversely, increasing excitability of RSC resulted in an increased firing rate in a subset of regions (red nodes in Figure 5.D), such as the right laterodorsal nucleus of the Thalamus (ldTh; Fig.5.D, right). Importantly, these changes do not involve all brain regions but a subset spanning a distributed and structured network.

Overall, these results demonstrate both focal and connectional diaschisis in silico. Specifically, they predict that modulating the excitability of a brain region produces a large-scale suppression of FC and local changes in firing rate activity in brain areas that are distant from the modulated region.

### 2.7 Inferring the mechanisms of network reconfiguration following focal region silencing

In the previous simulations, we just modified the excitability of one region without changing any other baseline parameter. However, local manipulations may lead to reorganizations in other brain regions, including their excitability (*η*) or their connections (*G*), as typically observed after brain lesions, e.g. after stroke [38] and epilepsy [39]. To account for possible structural and functional reorganizations, we used SBI to infer local and global parameters that best explain the data features before and after focal intervention in all mouse groups (Fig.6.A). Each violin plot in Figure 6.A represents the distribution of estimated parameters for several subsets of three mice. Namely, within each group (e.g., pre-silencing RSC), we randomly picked three mice and measured an array of average features (e.g., average FC). Then, we ran the SBI pipeline to estimate the working point where the simulations best match the average features of the mice subset. SBI predicts that focal region silencing induces both local and global alterations. Except for the CTRL group, the global coupling *G* was always decreased after focal silencing (Fig.6.A), indicating a decoupling between structure and function. This result is in line with the analysis in Figure 4.B where the correlation between structure and function decreased after RSC excitability was increased.

**Fig. 6.**
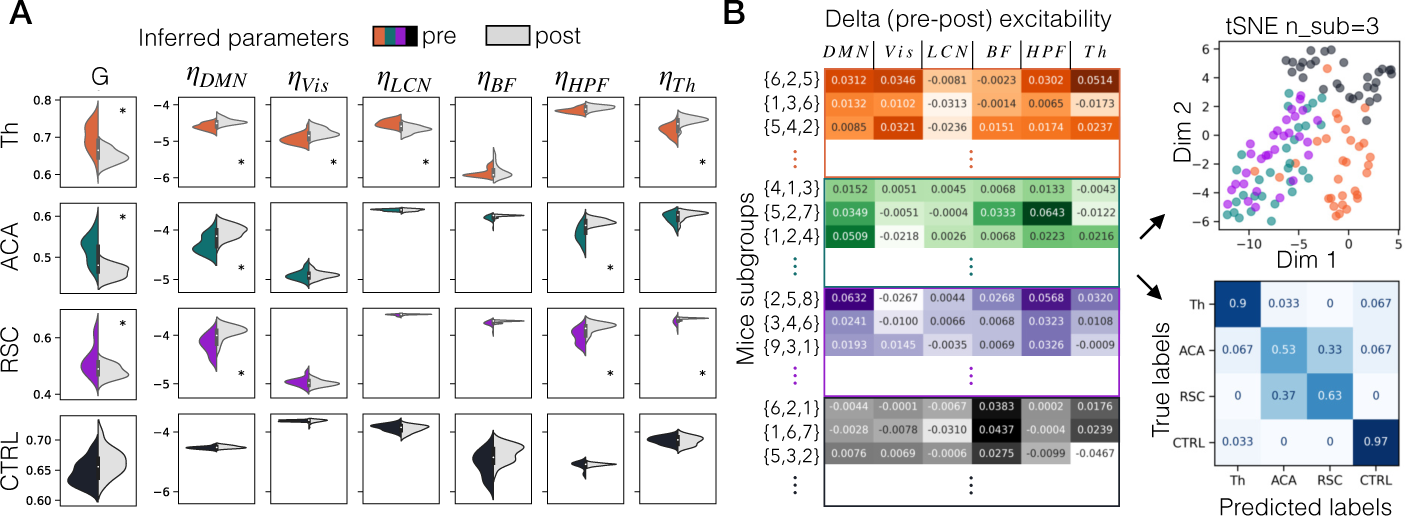
Local and global reparametrization and silencing classification: SBI was used to infer parameters in both the pre(colored) and post-silencing (gray) data for each mouse. The violin plots represent the distribution of inferred parameters. SBI predicts both local and global alterations resulting from focal region silencing; the global coupling G always decreases (except CTRL), whereas the local excitability parameter *η* can either decrease or increase. Significant shifts in the pre/post distributions are measured via the Wilcoxon test (*p* < 0.0001 (*)). B) The delta pre-post excitability was measured for different triplets of mice within each group. Then tSNE in a two-dimensional map is used to classify a given mouse triplet as belonging to a specific group. Based on the inferred delta excitabilities, the confusion matrix illustrates a high value of accuracy in the classification, particularly in Th, and CTRL groups.

Interestingly, the SBI analysis predicts that regional excitability can either decrease or increase after focal silencing (Fig.6.A). In particular, the RSC group presents a significant increase in the excitability of DMN, and in the sub-cortical hippocampal formation (HPF), and Th networks. This result extends the previous analyses (Figs.4-5), suggesting the possibility of non-trivial remote changes in excitability [38]. Importantly, the changes depend upon the type and location of the perturbation. In the ACA group, the excitability of the DMN and HPF were increased. In the Th group, excitability increased in DMN, Vis, and Th, whereas it decreased in the LCN.

### 2.8 Predicting the region where the manipulation occurred from experimental data

Based on the results of Figure 6.A, we hypothesized that silencing a specific site would result in a characteristic pattern of reconfiguration of regional excitability. If this is the case, we should be able to classify the Th, ACA, RSC, and CTRL groups based on the inferred changes in excitability. To test this hypothesis, we defined a six-dimensional array measuring the changes in RSN excitability (*η_pre_* − *η_post_*)*/*(*η_pre_* + *η_post_*) for each subset of mice in each group (Fig. 6.B). We used cross-validation to show that mice can be classified according to the effects of silencing on the network excitability. We then performed the cross-validation on 30 triplets of mice per group. In Figure 6.B (right), we show that mice can indeed be classified according to the changes in network excitability as estimated by SBI. Mice in the various groups were classified with 90% accuracy for Th, 53% for ACA, 63% for RSC, and 97% for CTRL. In other words, by providing the data before and after silencing from a non-specified group of mice, SBI enables us to accurately infer to which group the mice belong. The ACA and RSC groups were often mixed by the classification algorithm, due to the similarity of effects induced by silencing ACA and RSC, two strongly connected ROIs of the DMN.

## 3 Discussion

Both our empirical and theoretical results lend support to the concept of diaschisis [7]. Altering the excitability of even a single brain region can trigger widespread alterations in the network dynamics observable at the scale of the entire brain. This increases the complexity of data interpretation when utilizing perturbative approaches to investigate the mechanisms of brain function and dysfunction. For instance, if a behavioral change (Y) ensues following the targeted manipulation of a specific cell type (X) using optogenetic or chemo-genetic techniques, it might seem straightforward to infer a direct causative role of X in regulating behavior Y. However, this interpretation becomes less clear-cut when considering that the manipulation of X could trigger a broader reconfiguration of the entire dynamic landscape of the brain. In such a case, the observed behavioral outcome (Y) may indeed be a consequence of a systemic reorganization, with cell type X acting as a crucial, yet not unique, intermediary in this complex network. Our results are in line with previous results showing both local and non-local effects of brain lesions [7]. Experiments with rodent models of cortical lesions induced by photo thrombosis, which is evocative of cortex damage induced by stroke, reveal a perilesional reduction in gamma-aminobutyric acid receptor expression, paired with heightened contralesional NMDA receptor expression, suggesting excitation/inhibition imbalance in the two hemispheres [40]. In rats, in vivo chronic partial denervation of Schaffer collaterals projecting to hippocampal CA1 cells resulted in a homeostatic increase of CA1 excitability [41]. In humans, increased fMRI activity in motor and non-motor regions has been observed after stroke [42], especially in the contralesional areas [43–45]. These effects were also related to widespread cortical hyperexcitability [38, 46]. This reconfiguration could be explained by a mechanism of increased activation via decreased inhibition in long-range communication [47–49]. Also, stroke can induce changes in stimulus-evoked responses. For example, TMS stimulation revealed a lowered threshold for eliciting motor-evoked potentials and a higher evoked potential amplitude in stroke patients [50].

Our experiments confirm previous observations of decreased Functional Connectivity upon silencing of cortical hubs [18, 51]. Expanding this prior work, we show that structured and widespread reduction in long-range fMRI connectivity also characterizes subcortical thalamic lesions. Importantly, we show that the Functional Connectivity decrease stems from a compromised ability of the brain to sustain large-scale bursts of activity. While our results are consistent across manipulations and are statistically significant, the direction of the observed functional changes is in contrast with previous reports of paradoxical overconnectivity upon chemogenetic inhibition of the cortical and subcortical areas in mice and non-human primates [16, 17]. Several reasons may account for such discrepancies: different regions were targeted, the extension of the manipulated region, and the anesthetic combinations. More broadly, this apparent divergence reveals a complex and nuanced mechanism underlying the reconfiguration of Functional Connectivity.

To obtain mechanistic insights and make testable predictions, we employed The Virtual Mouse Brain software. This tool enables the simulation of electrophysiological activity throughout the entire brain, a feat not currently attainable experimentally. Furthermore, it allows the derivation of hemodynamic signals that can be compared to experimental BOLD signals. Using Simulation-Based Inference and seven free model parameters, we could reproduce whole-brain dynamics at baseline. To assess degeneracy [52], we would have to consider the excitability of each brain region as a free parameter. The computational cost prevented a complete analysis. Yet, it is quite remarkable to match experimental and theoretical results with only seven free parameters. The theoretical results led to the apparent paradoxical prediction that to explain the experimental results, inhibiting RSC cells with chemogenetics should increase local excitability. Remarkably, such non-trivial predictions were verified experimentally. This emphasizes the limitations of assessing the effects of focal manipulations without simultaneously considering their broader implications on global network dynamics. Thus, the optogenetic stimulation of parvalbumin cells can lead to their inhibition via a network effect [53]. In our study, we considered manipulation targeting regions considered as major hubs. It is possible that targeting other regions may produce different consequences at the whole brain scale. These effects may be predicted with whole-brain modeling.

Our model predicted that controlled inhibition leads to over-connectivity of hemodynamic signals, along with decreased firing rate, and increased (decreased) low (high)-frequency in neuroelectric signals. These results closely match the observations previously reported in [16, 17, 54]. A similar change in local field potential was also observed in intracranial electroencephalography recordings in humans where a focal lesion was performed by thermocoagulation [11]. Interestingly, the opposite effect is predicted for focal excitation, resulting in a decrease in long-range correlations, which better aligns with our experimental observations. The focal increase in excitation was paralleled by increased firing rate, decreased low frequency, and more robust high frequency in neuroelectric signals. This prediction of our model could be confirmed in future electrophysiology studies on the effects of increased focal excitability. Initial data, seem to support this prediction (Sastre D et al., *in preparation* 2023).

The insights gained from this study hold the potential for personalized predictions of the effects of local modulation on brain organization. Such knowledge could be harnessed for the development of targeted interventions [55], tailored neuromodulation therapies, aid in clinical decision-making, and advancements in brain-computer interface technologies. Importantly, we show that altering specific regions can lead to specific whole-brain dynamical signatures, which can be used as predictive biomarkers. This opens the way to new diagnostic possibilities in patients when there is no visible lesion on the MRI. Such mapping may reveal the locations of regions with altered function as compared to a control group.

In conclusion, our study underscores the intricate consequences of focal brain region perturbations, providing novel insights into brain function and the implications of localized modulation. By elucidating the interactions between neural scales, excitability changes, and remote effects, this study contributes to a deeper understanding of brain network dynamics and opens avenues for future research in both basic neuroscience and clinical applications.

## Author contributions

G.R, C.B, designed research; G.R., performed research; G.R., A.ZM., M.H., P.V., A.R., simulated the models and analyzed the results; HA.L., Z.L., P.Q., A.Gh., O.A., M.E., KH.C., AT.PB., A.V., performed the experiments; HA.L., Z.L., G.R., AT.PB., A.V., KH.C., preprocessed the experimental data; G.R, P.S., A.Go., V.J., C.B, organized the results; G.R., designed the figures; G.R., P.S., A.Go., V.J., C.B., wrote the paper. X.X contributed equally to this work as the last author. Correspondence should be addressed to Giovanni Rabuffo at or Christophe Bernard at.

## Data and materials availability

All data, code, and materials used in the analyses will be made available at a GitHub repository depending on the outcome of the editorial process.

## Supplementary information

### 5 Methods

In two independent experiments, we performed mouse brain imaging before and after region silencing (Fig.1,A). For region-level analysis, both datasets were registered onto a parcellation of the Allen Mouse brain consisting of *N* = 148 regions of interest (ROIs; Suppl. Fig.S2).

#### 5.1 Lesion datasets

All experiments were conducted in accordance with Inserm and Aix-Marseille Universitè Ethical Committee guidelines. The protocol was approved by the French MESRI under authorization #30794-2021033018524869v4. All surgical procedures were conducted under anesthesia, with the utmost care taken to minimize distress and ensure the overall well-being of the animals from their arrival to their death. All the animals were housed in large cages of 4 to 6 individuals with enrichment, food, and water at libitum, in a ventilated cabinet providing a controlled environment (temperature: 22 ± 1°C; 12 h light/dark schedule with lights off at 8:00 P.M.; hygrometry: 55%; ventilation: 15 − 20 vol/h). The ‘lesion’ dataset was obtained from 9 C57BL6 mice (Charles River, 3 males, and 6 females, 19 − 26g) with irreversible lesions in thalamic nuclei induced under ketamine + xylazine (100mg/kg, 10mg/kg, i.p.) or isoflurane (2.5%) by injections of sterilized NMDA 0.1M prepared in 0.12M sodium phosphate buffer, pH 7.2 − 7.4 (PB) using a Nanoject III (Drummond Scientific) with a slow injection rate (0.03*µ*L/min) over 10 min. The glass pipette was retained in place for another 10 min and then slowly retracted. After injection, the craniotomy was closed using Dura-Gel (Cambridge Neurotech) and Metabond (Parnell), and the skin was sutured. BOLD rs-fMRI acquisitions were performed before and 6 weeks after surgery under light anesthesia (0.6% isoflurane) and sedation (s.c injection of a single dose of medetomidine 0.13 mg/kg) on a 7T Pharmascan Bruker equipped with a cryoprobe using 2D GEEPI (spatial resolution of 0.167 × 0.167 × 0.40mm^3^; echo time of 16.30 ms; tBOLD Repetition Time TR= 1.75 s, flip angle of 30°, 15 min, 512 repetitions). Data processing (FSL software) included motion correction, regression of the volumes affected by motion and by sudden changes in BOLD signal intensity, skull stripping, bias field correction, slice timing correction, grand-mean intensity normalization, band pass filtering (0.01 − 0.1Hz), registration to Allen Template using linear and nonlinear transformations and spatial smoothing with a Gaussian Kernel of 4 mm (= 0.4mm in original resolution)[56, 57]. 3 mice were excluded since they did not present evident lesions from histology experiments, or because artifacts were present in the fMRI activity.

At the end of the recording, mice were deeply anesthetized by intraperitoneal injection (i.p.) of Ketamin (100mg/kg) / Xylazine (10mg/kg) solution then by a lethal i.p. injection of Euthasol (150mg/kg) and transcardially perfused with100 ml of 4% paraformaldehyde (PFA) prepared in PB. After perfusion, the brains were removed from the skull, post-fixed in the same fixative for 1h at room temperature (RT), rinsed in PB, and cryoprotected in a solution of 20% sucrose in PB overnight at 4°C, quickly frozen on dry ice and sectioned coronally at 40*µ*m with a cryostat. The sections were rinsed in PB, collected sequentially in tubes containing an ethylene glycol-based cryoprotective solution, and stored at −20°C until histological processing. Every 2 sections, was stained with cresyl violet to determine the general histological characteristics of the tissue along the rostrocaudal axis of the brain and the extent of the lesion.

#### 5.2 DREADDs datasets

The ‘DREADDs’ dataset was obtained by DREADD inhibition of the retrosplenial cortex (RSC; N = 14) or anterior cingulate area (ACA; N = 7) on C57Bl/6J male mice, which are generally considered functional hubs in the mouse brain [18, 33]. Another control group (N = 8) was also included in which a control virus was injected into RSC. As RSC is a very elongated area, we targeted the posterior section based on our preliminary study which shows stronger functional connectivity within DMN. Surgeries were performed at least one month before fMRI. The animal (10 – 16 weeks of age) was anesthetized with 1.5 − 2% isoflurane during surgery. Enrofloxacin (6 mg/kg) and carprofen (5 mg/kg) were injected subcutaneously to prevent infection and relieve pain and inflammation, respectively. Body temperature was maintained at 37 °C with a heating pad. In ACA and RSC groups, inhibitory DREADD virus AAV2/1 pSyn-hM4D(Gi)-T2A-mScarlet was bilaterally injected into ACA or RSC with 0.25 µL for each side. In the CTRL group, control virus AAV DJ/8 pAM-tdTomato was bilaterally injected into RSC with 0.25 µL for each side. Both DREADD and control viruses were obtained from Sah lab, Queensland Brain Institute, The University of Queensland, QLD, Australia. The relative coordinates to Bregma of ACA and RSC based on Paxions and Franklin Mouse Brain atlas, fifth edition are:

- Anterior cingulate area (ACA). ML: ±0.3mm; AP: +0.4mm; DV: -0.9mm.
- Retrosplenial cortex (RSC). ML: ±0.4mm; AP: -2.7mm; DV: -0.6mm.

The virus was injected using a Nanoject III (Drummond Scientific) with a slow injection rate (0.03 *µ*L/min) over 10 min. The glass pipette was retained in place for another 6 min and then slowly retracted. After injection, the wound was closed using Vetbond (3M) and sutured. Enrofloxacin and carprofen were administered for another two days. Animals were kept in their home cage (group housing of 2-4 animals per cage) for 4 weeks to recover and to allow expression of the virus before MRI. MRI was conducted on a 9.4T system (BioSpec 94*/*30, Bruker BioSpin MRI GmbH). Animals were initially anesthetized using 3% isoflurane in a 2:1 air and oxygen mixture. After being secured in an MRI-compatible holder using custom-made tooth and ear bars, a bolus of medetomidine was delivered via an i.p. catheter (0.05−0.1 mg/kg), and the isoflurane level was progressively reduced to 0.25 − 0.5% over 10 min, after which sedation was maintained by a constant i.p. infusion of medetomidine (0.1mg/kg/h). Key physiological parameters, including arterial oxygenation saturation (SpO2), rectal temperature, heart rate, and respiratory rate, were measured by an MRI-compatible monitoring system (SAII Inc). Body temperature was maintained at 36.5°C with a heated water bath. About 45min after the bolus injection of medetomidine, the first 10 min resting-state fMRI data was acquired using multiband EPI as a pre-DREADD scan (TR/TE = 300*/*15 ms, thickness = 0.5 mm, gap = 0.1 mm, 16 axial slices covering the whole cerebrum with in-plane resolution of 0.3 × 0.3 mm^2^) [58]. After this, 1mg/kg water-soluble clozapine N-oxide (CNO dihydrochloride; cat #6329, Tocris Bioscience) was slowly injected via a pre-implanted catheter (i.p.) in 1 min. 30 min after the CNO injection, another 10 min resting-state fMRI was performed as a post-DREADD scan. The data was preprocessed by motion correction, distortion correction, nuisance removal [59], bandpass filter (0.010.3Hz), transformed to ABA 200*µ*m template with 10× resolution (2mm) and then smoothed by 6mm Gaussian (= 0.6mm in original resolution).

#### 5.3 Empiric structural connectivity

The connectome used in the simulations was obtained by employing The Virtual Mouse Brain pipeline (TVMB; [20]) on tracer experiments performed at the Allen Institute ([28]). In these experiments, adult male C57Bl/6J mice were locally injected with a recombinant adeno-associated virus expressing the EGFP anterograde tracer. The tracer migration signal was captured using a serial two-photon tomography system, providing information about the axonal projections originating from the injected sites. Injection density was defined as the ratio of infected pixels to the total number of pixels in the source region. Similarly, projection density was defined in each target region as the ratio of detected pixels following injections in the source region to the total number of pixels in the target region. By averaging injection experiments conducted in the right brain areas and targeting regions in both the ipsilateral and contralateral hemispheres, a tracer-based connectome was constructed. Structural connection strength between a source region *n* and a target region *m* (referred to as the link *nm*) was determined by calculating the ratio of the projection density at region *m* to the injection density at region *n*. The resulting tracer connectome displayed normalized links ranging between 0 and 1.

#### 5.4 Neural mass model

In whole-brain simulations, the activity of each ROI is modeled by suitable dynamical equations describing the mean-field behavior of an isolated neural mass. In this work, we used a neural mass model (NMM) which describes the co-evolution of mean firing rate *r* and mean membrane potential *V* of a large population of all-to-all coupled quadratic integrate and fire (QIF) neurons [60]. The local region *n* dynamics is described by the following NMM equations

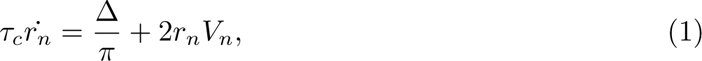

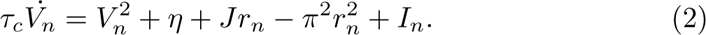

Here we used default parameters *η* = −4.6 for the average excitability of the QIF neurons, Δ = 0.7 for the spread of the neuronal excitability distribution in the population, *J* = 14.5 for the synaptic weight across coupled QIF neurons, and *τ_c_* = 1 for the characteristic time. In this configuration, the decoupled neural mass model is in a bistable regime [30], with a UP state corresponding to a stable focus and a DOWN state corresponding to a fixed point. The input *I_n_* is additive in the membrane potential equation.

#### 5.5 Virtual brain simulations

Connecting the NMM equations of each brain region through the Allen structural connectome defines a system of coupled equations that models the mouse brain dynamics. When a region *n* is coupled to other brain regions through the structural connectivity matrix *W_nm_*, the total input to region *n* is

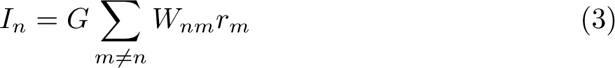

The global coupling parameter *G* regulates the influence of the connectome over the local dynamics. When *G* = 0 the regions are decoupled and the brain regions evolve according to the local NMM dynamics. The more *G* increases, the more the brain regions’ dynamics are intertwined. In this work, we do not account for signal transmission delays, and we do not simulate excitatory versus inhibitory population dynamics (see limitations in Discussion. The solution of the coupled system was obtained by numerical integration using the TVMB platform [19, 20], and corresponds to a simulated dataset describing the evolution of the 2*N* variables (*r_n_*(*t*)*, V_n_*(*t*)). The sampling rate of these neuroelectric variables is set to 1000Hz. The simulated BOLD activity *B_n_*(*t*) in each region is derived by filtering the firing rate *r_n_*(*t*) through the Balloon-Windkessel model [61], modeling the hemodynamic response function of the BOLD signal. The parameters were tuned to match realistic hemodynamic timescales for the mouse (Suppl.Fig.5). We use a repetition time of 2 seconds so that the BOLD rate is 0.5 Hz.

#### 5.6 Simulation-based inference (SBI)

Bayes’ rule describes how the prior (background information before seeing the data) is combined with likelihood (available information in the data) to form the posterior distribution for making probabilistic inferences and predictions [62]. Mathematically speaking, the posterior distribution of parameters *θ* given the model *m* is *p*(*θ* | *y, m*) ∝ *p*(*y* | *θ, m*)*p*(*θ* | *m*), which *p*(*y* | *θ, m*) represents the likelihood and *p*(*θ* | *m*) is the prior information [63]. For many high-dimensional and nonlinear dynamical models, such as The Virtual Mouse Brain (TVMB) used in this study, the calculation of the likelihood function becomes intractable [35], and standard methods, such as Markov chain Monte Carlo sampling, cannot be applied directly to carry out Bayesian inference. Hence, likelihood approximation algorithms such as Simulation-based Inference (SBI; [21]) are required to efficiently estimate the posterior of the unknown parameters. To perform SBI without sensitivity to a distance measure for model evaluation [64], deep neural density estimators can be trained on a set of forward simulations (generated with random parameters drawn from a prior), to learn the likelihood function. In particular, using invertible transformations through Masked Autoregressive Flow (MAF; [65]) in a class of SBI methods (the so-called Sequential neural posterior estimators, SNPE; [66]), we are able to readily estimate the approximated posterior *q_φ_*(*θ* | *y, m*) with learnable parameters *φ*, so that for the observed data *y_obs_*: *q_φ_*(*θ* | *y_obs_, m*) ≃ *p*(*θ* | *y_obs_, m*).

In this study, we ran a MAF in one round of SNPE, with 5 autoregressive layers, each with two hidden layers of 50 units, as implemented in the PyTorch-based SBI package [67]. To estimate the joint posterior distribution between parameters *G*, and *η*, we performed 10^5^ simulations with a uniform prior as *G* ∈ U (0, 1.1) and *η_i_* ∈ U (−6, −3.5).For the training of the neural network, we utilized features extracted from the BOLD signal. These features were related to static and dynamic network measures: the Functional Connectivity (FC) was measured as the pairwise Pearson’s correlation among ROIs-level signals; the dynamic Functional Connectivity (dFC) is a (time× time) matrix defined as the Pearcon’s correlation among pairs of instantaneous co-activation patterns *E_ij_*(*t_A_*) · *E_ij_*(*t_B_*) (vertical slices of the green block in Figure 2.A) at times *t_A_* and *t_B_*. The features used in the training included the fluidity (i.e., the variance of dFC), the overall sum of FC and dFC matrices, the average value of FC homotopic, and the variance of the masked dFC matrix over six sub-networks: DMN (Default Mode Network), Vis-Aud (Visual-Auditory), LCN (Limbic-Cortical Network), BF (Basal Forebrain), HPF (High-Processing Frontal), and Thalamus.

#### 5.7 Surrogate data

In order to compare our results in 2 a with the null hypotheses of random evolution and of inter-regional stationarity of the data, we build phase-randomized surrogates. According to [68] the phase randomized surrogate is obtained by adding a uniformly distributed random phase to each frequency of the Fourier-transformed signal, and subsequently retrieving the phase-randomized signals by inverse Fourier transform. Importantly, the phase shift is different in every frequency but can be applied uniformly to all the brain regions (cross-spectrum preserved) or separately in every region (cross-spectrum not preserved). Only in the first case, the static *FC* is preserved. The coherent fluctuations around the stationary *FC*, however, are destroyed. For more details see also [69].

**Fig. S1.**
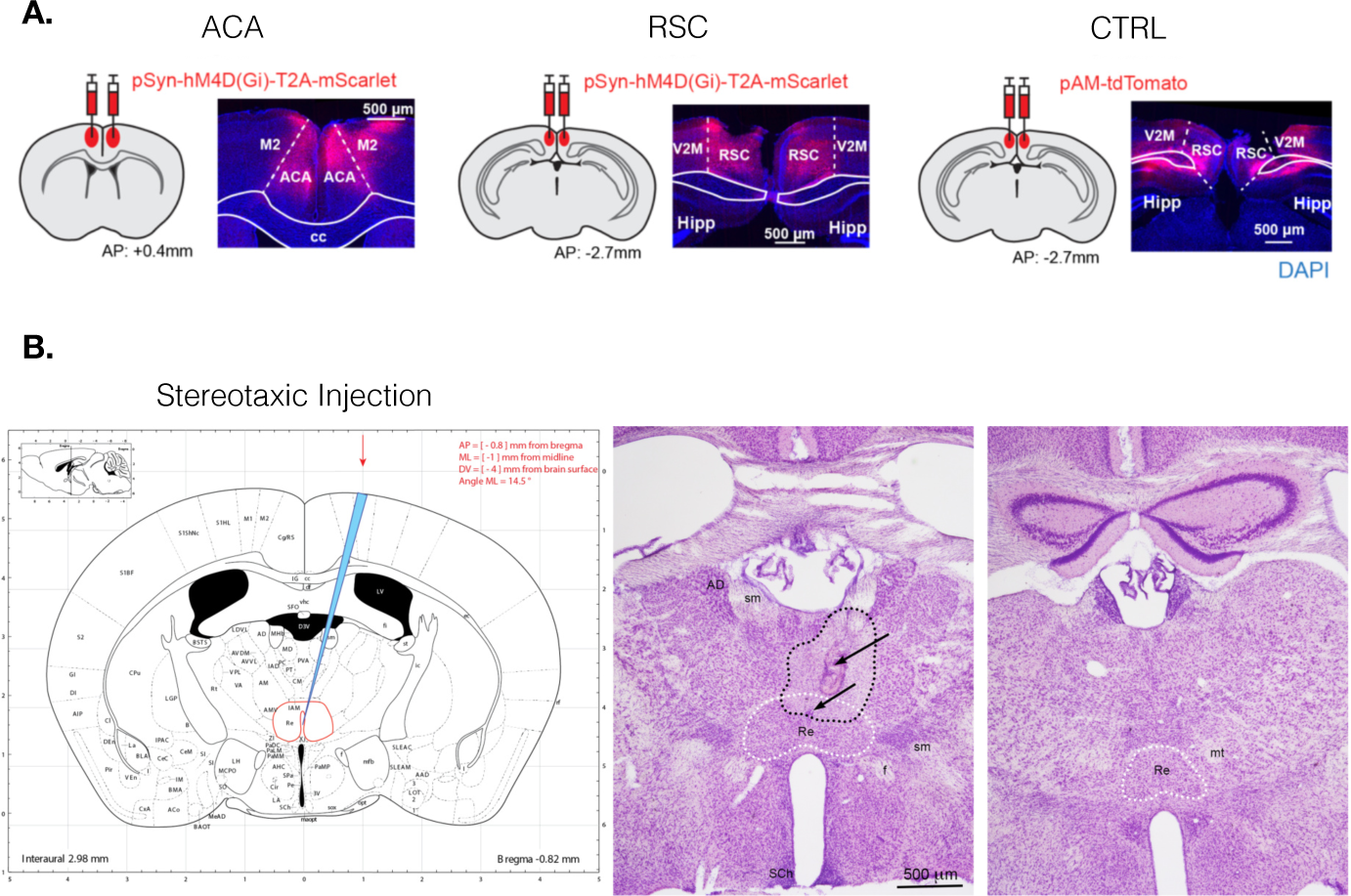
DREADD virus expression and thalamic lesions.: (A) DREADD expression in ACA, RSC, and CTRL groups. In each subgraph, the left image shows the injection location of the inhibitory DREADD (AAV-pSyn-hM4D(Gi)-T2A-mScarlet) or control (pAM-tdTomato) virus, and the right image shows the fluorescence imaging together with the DAPI staining (blue). cc, corpus callosum; ACA, anterior cingulate area; Hipp, hippocampus; M2, secondary motor cortex; RSC, retrosplenial cortex; V2M, the medial secondary visual cortex. (B) Example of stereotaxic injection of NMDA leading to lesion of central thalamic nuclei as illustrated on mouse brain sections processed for cresyl violet histological staining.

**Fig. S2.**
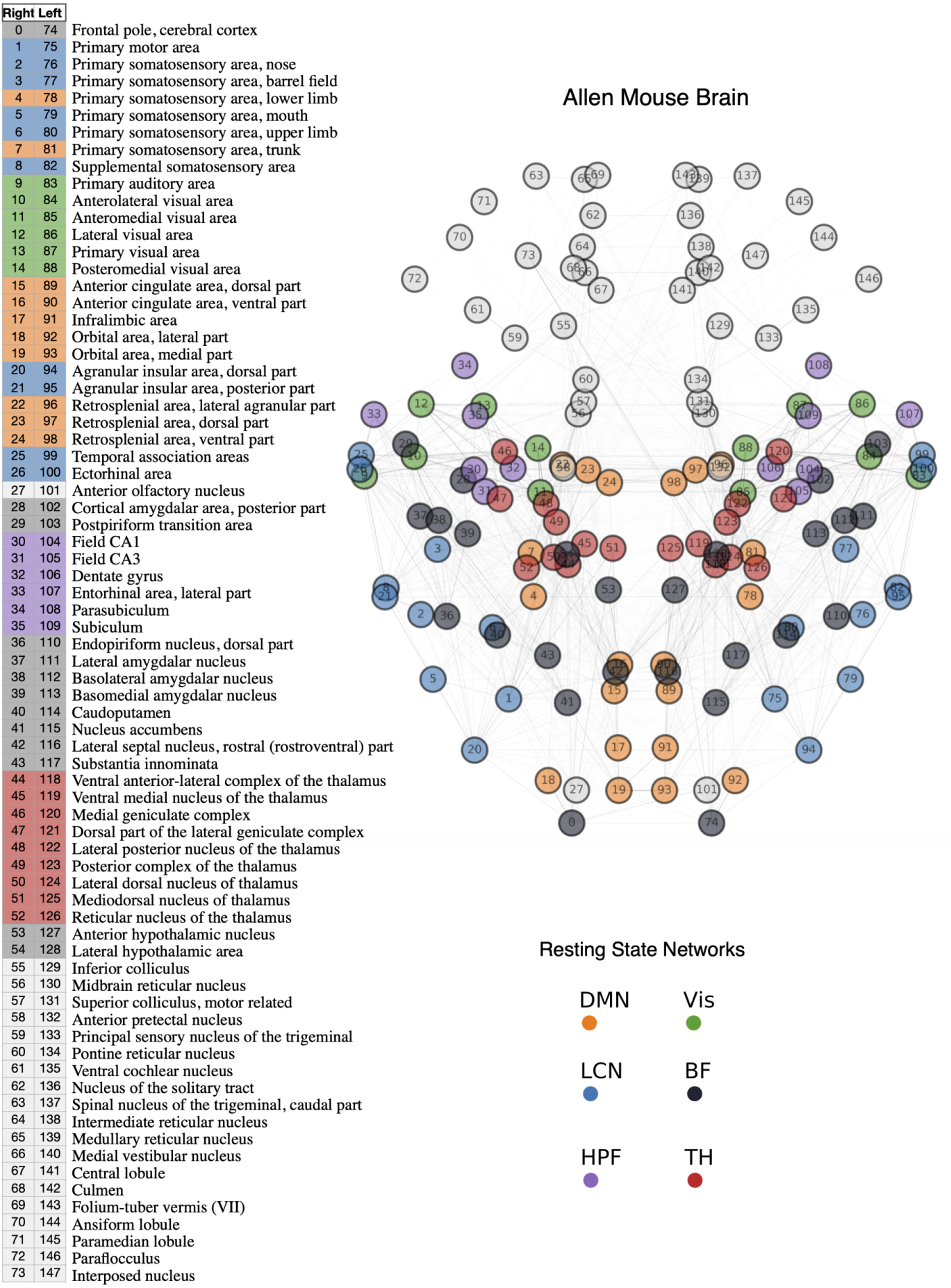
Allen Mouse Brain parcellation: This parcellation was obtained by processing the Allen Mouse data [28] as in [20].

**Fig. S3.**
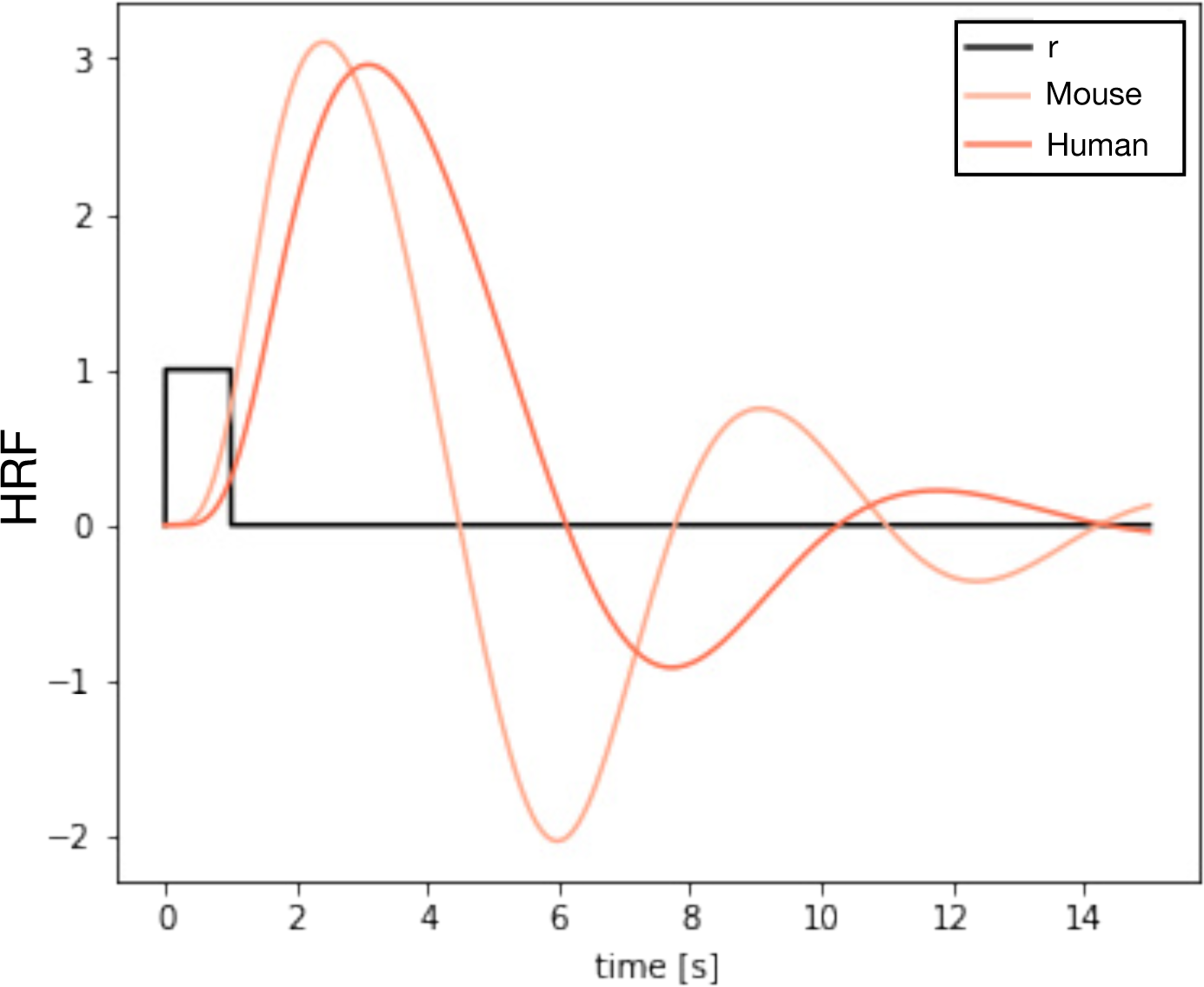
BOLD Hemodynamic Response Function: For a pulse of the firing rate *r* the TVB default hemodynamic response function (HRF) for the human is shown. We modified this kernel to match more realistic timescales observed in rodents [70]. The HRF is used to transform simulated firing rates into simulated BOLD data.

#### 5.8 Appendix A: ROIs-level FC analysis

We reconstructed ROIs-level BOLD signals according to an Allen parcellation with *N* = 148 ROIs. We computed FC correlation matrices Supp.Fig.3.A and we organized brain regions in mouse resting-state networks identified according to a common classification (as in Supp.Fig.2): Default Mode Network (DMN), Lateral Cortical Network (LCN), Visual Network (Vis), Basal-Frontal Network (BF), Hippocampal Formation (HPF), and Thalamus (TH). Averaging FC blocks (i.e., edges within black lines in Supp.Fig.3.A), we measured the average FC within and across resting state networks. Comparing pre- and post-silencing for RSC group shows that all inter- and intra-network FC connections decreased after the chemogenetic silencing (Supp.Fig.3.B, matrix numbers mark the result of Wilcoxon test, up/down arrow marks increased/decreased FC, color indicates p-value). The RSC group showed the largest FC decrease. The same procedure applied to ACA group showed a significant decrease only in the FC across LCN and DMN connections, whereas in the Th group, only the correlations across LCN and Vis decreased significantly (Supp.Fig.3.B). The FC decrease for each RSC mouse was measured by Δ*FC* = *FC_post_* − *FC_pre_* Supp.Fig.3.C and it was significant across most brain networks when compared to the CTRL group (Supp.Fig.3.D; Mann-Whitney U test, FDR corrected). Thus, also at the (ROIs-reconstructed) edge level, the values of the observed differences in the RSC group are greater than those in the CTRL group.

The deltas *FC_post_* − *FC_pre_* were averaged across subjects within the corresponding group (Supp.Fig.3.C). To provide a topography of the differences we selected all edges in the RSC group that showed greater differences than the highest difference observed in the CTRL group (Supp.Fig.3.C). The resulting network is shown in Supp.Fig.3.E. For ACA and Th groups, we show the same number of edges with the highest differences (Supp.Fig.3.E, center, and right, respectively). While silencing RSC and ACA resulted in a suppression of functional networks connected to ACA and RSC, the Thalamic lesions mainly decoupled the interhemispheric connections, with a decrease in correlation between homotopical regions and weaker effects on Thalamic nuclei.

**Fig. S4.**
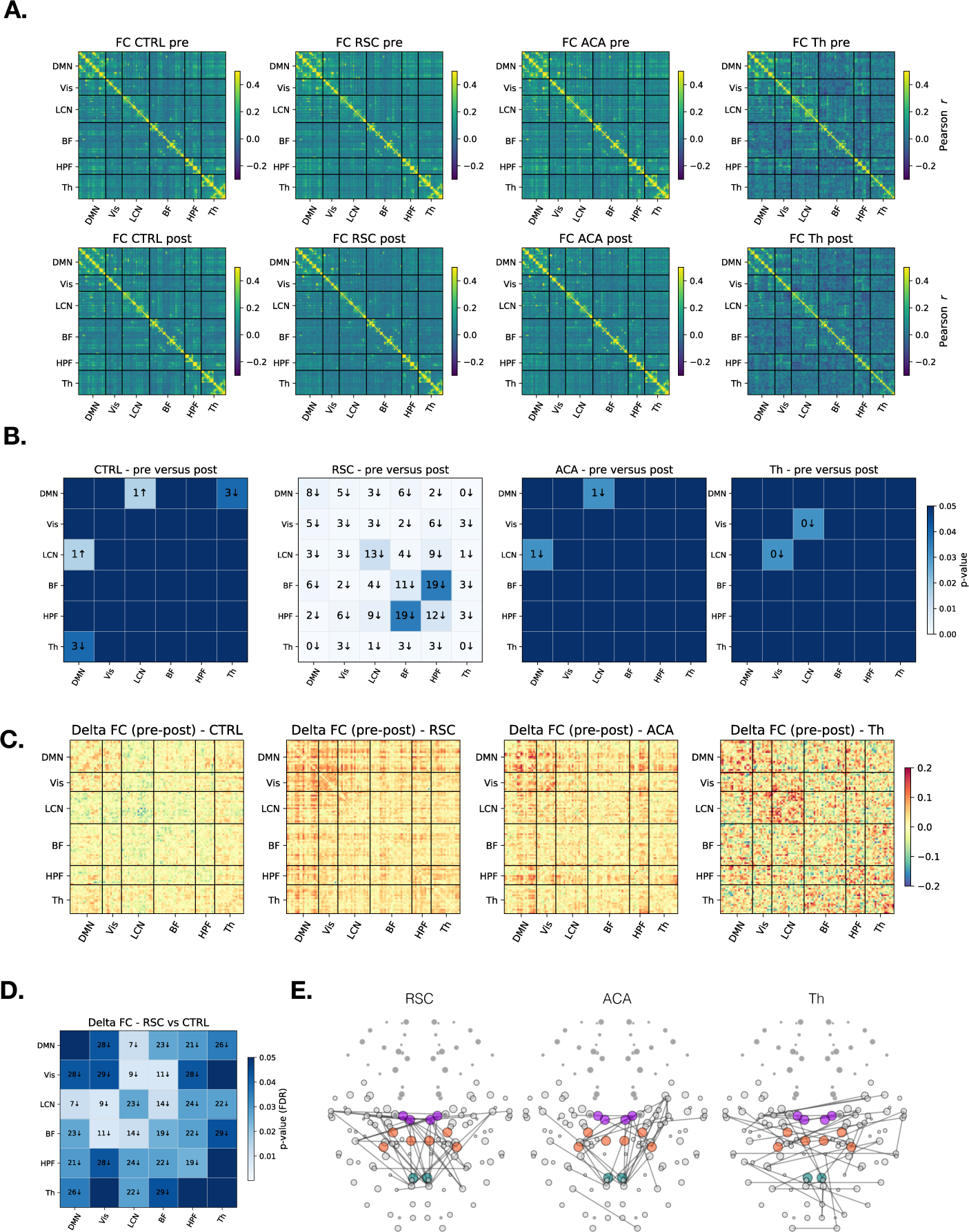
Region-level Functional Connectivity changes after focal silencing: A) FC at the ROI-level was measured in all mice for pre- and post-silencing trials. ROIs were sorted according to resting state network (RSN) classification (see Methods). B) FC was averaged within and across RSNs (blocks delimited by black lines in panel A), and the statistical significance of the FC decrease was assessed in the RSC group by comparing pre- and post-FC values. B) Each matrix shows the significance of all intra- and inter-RSN changes. In each entry, the numbers indicate the value of the Wilcoxon test, the color indicates the p-value, and the arrows (all pointing down) indicate that correlations within or across RSN always decreased after the focal silencing of RSC. C) Average delta (post-pre) FC in the CTRL, RSC, ACA, and Th groups. D) The delta FC in the RSC mice was significant when compared to the delta FC of the CTRL mice (Mann-Whitney U test, FDR corrected; colors and labels as in panel (B). E) The strongest correlation decreases are plotted for the RSC, ACA, and Th groups.

#### 5.9 Appendix B: SBI results from a different TVB parametrization

We use SBI to infer the configuration of local and global parameters (*η_ACA_, η_RSC_, η_T h_, G*) that best explain the data features in the pre and post-silencing cases (see Methods). SBI predicts that focal region silencing induces local excitability changes and global network segregation, as measured by a decrease in the global coupling Supp.Fig.4.A, left. As expected, these changes were not significant in the CTRL group, as opposed to the RSC group. In particular, in Supp.Fig.4.A, right, we show that RSC, ACA, and Thalamus all have higher excitability after direct silencing, i.e., by local intervention. On the one hand, when the thalamus is lesioned, it is not predicted that the excitability of cortical hubs will change significantly. On the other hand, following either ACA or RSC silencing, the model infers decreased thalamic excitability *η_T h_* and increased cortical excitabilities *η_ACA_*and *η_RSC_*. In other words, chemogenetic silencing is predicted to induce nonlocal changes in regions that are not directly silenced. The ACA and the RSC are strongly connected regions, as both are part of the DMN, and they show concurrent increases when either is shut down. The thalamus, on the contrary, is inferred to show decreased excitability after the chemogenetic silencing of these cortical hubs, which might be interpreted as a compensatory mechanism. To check for the robustness of our results, we have also used the model to infer the parameters that were best predictive of average features across subsets of mice (as opposed to fitting each mouse individually). As evident in Supp.Fig.4, we show that, as average features are evaluated over a growing number of mice, the results quickly converge to an even sharper distinction between the pre-and post-conditions (with the exception of the control group), in the same direction as those observed at the single-mouse level.

In conclusion, in this section, we have used probabilistic machine learning and mechanistic models to infer the most plausible local, microscopic mechanisms through which local lesions affect large-scale patterns of activities. We conclude that a decrease in network integration and an increase of excitability in the silenced regions, accompanied by nonlocal excitability changes, are possible candidates for causes of impairment.

**Fig. S5.**
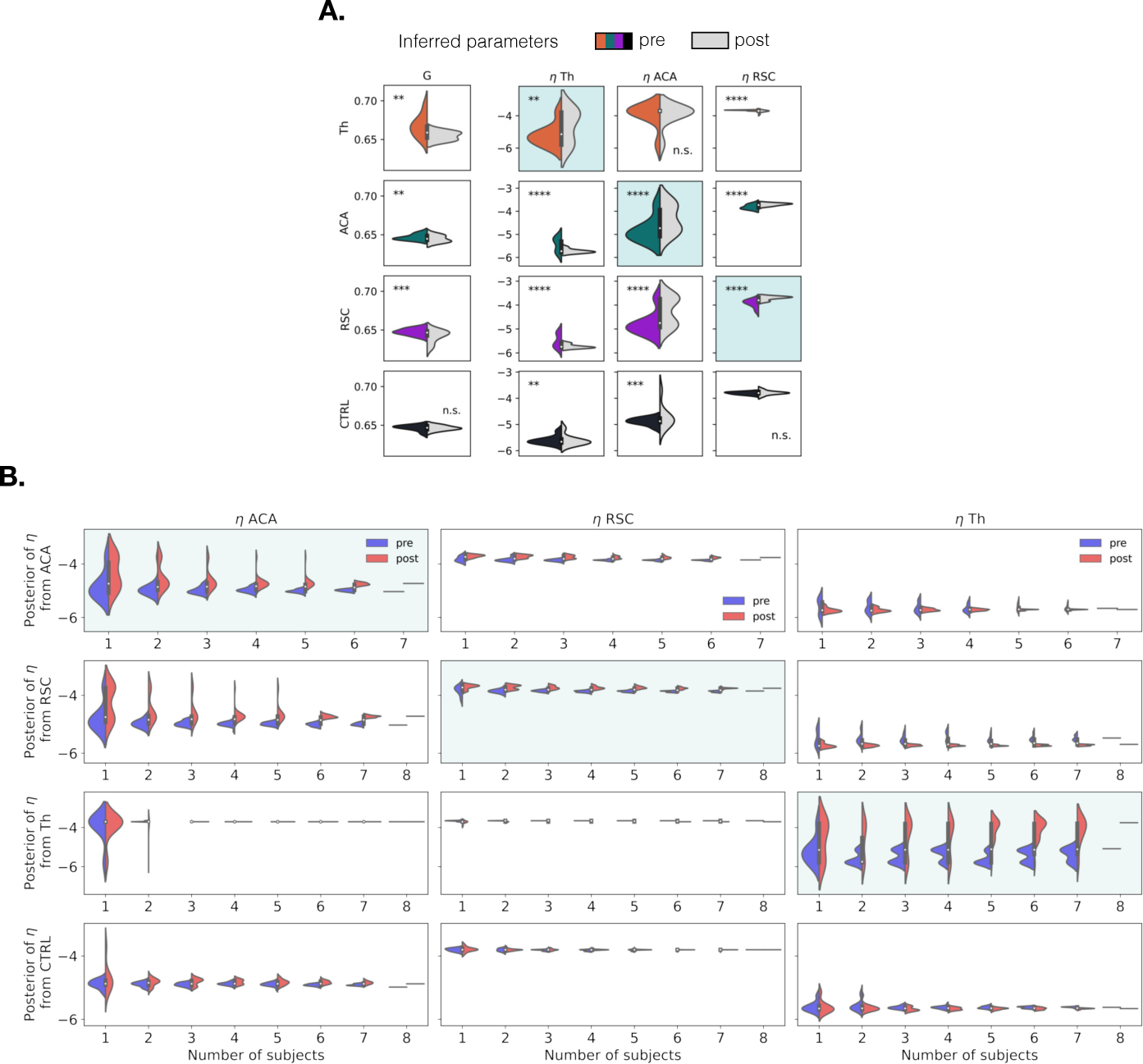
SBI results from a different TVB parametrization. (A) The estimated posterior distribution of the global coupling parameter *G* and the excitability parameter *η* in ACA, RSC, and Th groups, for pre-lesioning and post-lesioning (*p* < 0.01 (**), *p* < 0.001 (***), *p* < 0.0001 (****)), obtained by training a probabilistic deep learning algorithm, the so-called MAF (see Methods), on the summary statistics of simulations. Results indicate a significant increase in the corresponding regions by lesioning. (B) The estimated posterior distribution of the excitability parameter *η* in ACA, RSC, and Th groups, for pre-lesioning and post-lesioning. Here, the summary statistics are averaged over different subsets of mice.

